# Diffusion Basis Spectrum Imaging of White Matter in Schizophrenia and Bipolar Disorder

**DOI:** 10.1101/2024.07.07.602402

**Authors:** Daniel Mamah, Aakash Patel, ShingShiun Chen, Yong Wang, Qing Wang

## Abstract

**Background:** Multiple studies point to the role of neuroinflammation in the pathophysiology of schizophrenia (SCZ), however, there have been few *in vivo* tools for imaging brain inflammation. Diffusion basis spectrum imaging (DBSI) is an advanced diffusion-based MRI method developed to quantitatively assess microstructural alternations relating to neuroinflammation, axonal fiber, and other white matter (WM) pathologies.

**Methods:** We acquired one-hour-long high-directional diffusion MRI data from young control (CON, *n* =27), schizophrenia (SCZ, *n* =21), and bipolar disorder (BPD, *n* =21) participants aged 18-30. We applied Tract-based Spatial Statistics (TBSS) to allow whole-brain WM analyses and compare DBSI-derived isotropic and anisotropic diffusion measures between groups. Clinical relationships of DBSI metrics with clinical symptoms were assessed across SCZ and control participants.

**Results:** In SCZ participants, we found a generalized increase in DBSI-derived cellularity (a putative marker of neuroinflammation), a decrease in restricted fiber fraction (a putative marker of apparent axonal density), and an increase in extra-axonal water (a putative marker of vasogenic edema) across several WM tracts. There were only minimal WM abnormalities noted in BPD, mainly in regions of the corpus callosum (increase in DTI-derived RD and extra-axonal water). DBSI metrics showed significant partial correlations with psychosis and mood symptoms across groups.

**Conclusion:** Our findings suggest that SCZ involves generalized white matter neuroinflammation, decreased fiber density, and demyelination, which is not seen in bipolar disorder. Larger studies are needed to identify medication-related effects. DBSI metrics could help identify high-risk groups requiring early interventions to prevent the onset of psychosis and improve outcomes.

## Introduction

Schizophrenia (SCZ) is a chronically disabling psychiatric disorder afflicting 0.5-1% of the population. The pathophysiology of the disorder is unclear and complicated by the heterogeneity of clinical, postmortem, and imaging findings. However, SCZ has been associated with reductions in cortical thickness, as well as cortical surface area, primarily in frontal and temporal regions.^1^ Explaining the underlying causes of cortical changes, neuropathological studies have shown increased neuronal packing density in the cortex, due to both decreased neuropil and soma size.^2^ Dendrites are shorter and less branched and spine density is also lower in those with SCZ compared to healthy individuals,^3^ which is thought to underlie the neuronal connectivity abnormalities reported in SCZ patients.^4^ Additionally, fewer oligodendrocytes in various brain regions have been found in SCZ further contributing to neuronal dysconnectivity in the disorder, and white matter deficits found in brain imaging studies.^2^

White matter volumes are often reduced in SCZ patients, but generally to a lesser extent than cortical volumes.^5^ Diffusion imaging studies in SCZ typically report lower fractional anisotropy (FA) in one or more white matter tracts, however, the specificity of significant abnormalities has been heterogeneous across studies.^6–8^ The large, multi-site ENIGMA study involving 4322 individuals found widespread FA reduction in SCZ, with the anterior corona radiata (*d*=0.40) and corpus callosum (*d*=0.39) showing the greatest effects.^9^ Myelin deficits are thought to play a major role in observed diffusion imaging abnormalities, considering that increased radial diffusivity is usually seen in SCZ,^10^ and dysregulation of myelin-associated gene expression, reductions in oligodendrocyte numbers, and marked abnormalities in the ultrastructure of myelin sheaths have also been reported.^11^

Multiple lines of evidence have attributed neuroinflammation to SCZ,^12^ which appears to precede the onset of illness.^13^ Peripheral inflammatory markers (e.g., IL-1β, IL-6, and TNF-α) are often seen in individuals with SCZ,^14,15^ although diagnostic specificity remains to be established and is also seen in other psychiatric disorders.^15^ Robust evidence has also been reported for increased proinflammatory cytokines in the prodromal period,^16^ and prenatal exposure to specific infections has been associated with future SCZ development.^17^ Furthermore, results from genetic studies support the role of immune dysregulation in the pathophysiology of SCZ – a large multi-site study found that the most significantly associated SCZ genome-wide association study (GWAS) locus lies within the major histocompatibility complex (MHC) locus on chromosome 6, which contains genes that function in the acquired immune system.^18^ Finally, positron emission tomography (PET) studies done in SCZ patients, using a variety of tracers, most commonly targeting the 18 kDa translocator protein, TSPO, have shown increased binding in the disorder.^12^ While PET studies can measure neuroinflammatory proteins at low concentrations, obstacles to reliable measurement remain, such as high nonspecific binding and sensitivity to a genetic polymorphism that affects the binding affinity of early TSPO radioligands.^19^ Other promising neuroinflammation-targeted ligands are currently in development.^19^ Outside of PET, few non-invasive alternatives for investigating neuroinflammation in the brain have been proposed.^20^ In contrast to PET, such methods could facilitate longitudinal investigations, particularly in younger populations, and potentially have future clinical implications in monitoring treatment effects.

Diffusion Basis Spectrum Imaging (DBSI) is a novel multi-parametric diffusion imaging post-processing methodology that models the diffusion-weighted MRI signals as a linear combination of multiple tensors describing both the discrete anisotropic content (axonal fibers) and an isotropic diffusion spectrum component encompassing the full range of diffusivities.^21,22^ DBSI acquires water diffusion signals for each imaging voxel and then uses a pattern-matching algorithm to find the minimal number of dictionary entries that best represent the acquired water diffusion signals and their relative proportions. Each identified dictionary entry represents one type of microstructure in the image voxel.^21,22^ This modeling was developed to quantify intra- and extracellular water permitting the discrimination of vasogenic edema, increased cellularity, axonal injury/loss, and demyelination (see **Figure 1**).^21,22^ Previous studies using DBSI have been done in Alzheimer’s disease where it showed increased cellular diffusivity in both preclinical and early symptomatic phases of Alzheimer’s disease consistent with white matter inflammation,^23,24^ despite traditional diffusion imaging abnormalities only identified in the symptomatic phase.^23^ DBSI has also been used in multiple sclerosis,^25^ cervical spondylotic myelopathy,^26^ obesity,^27^ and HIV,^28^ but has not been used to investigate individuals with SCZ or psychosis.

**Figure 1:**
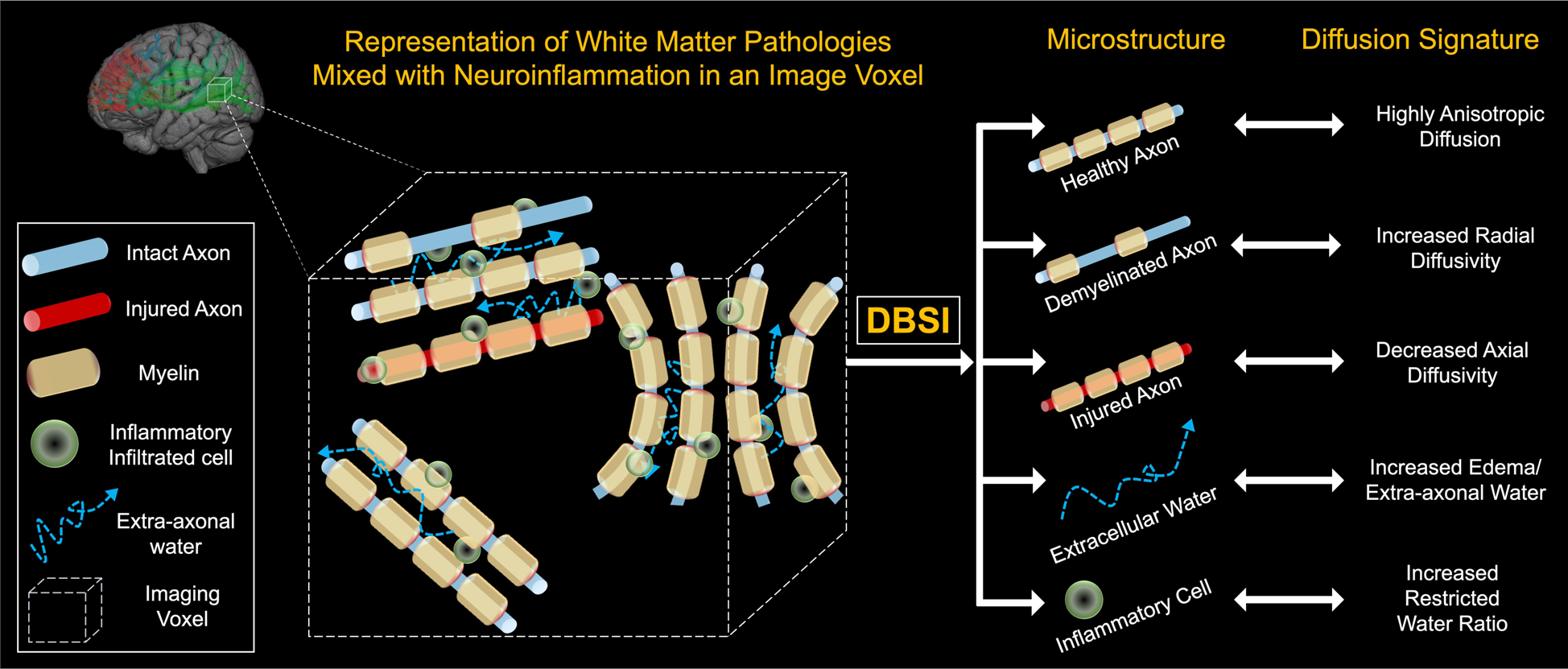
Schematic demonstration of Diffusion Basis Spectrum Imaging (DBSI). A data-driven multipole tensor model is employed by DBSI to model the white matter pathophysiology. Based on the diffusion signatures, DBSI can separate various sub-voxel compartments within the imaging voxel to generate pathologically and physiologically specific imaging biomarkers such as cellular diffusion fraction, axonal water fraction, fiber specific axial and radial diffusivities. In addition, the method can estimate axonal fiber fraction which correlates with axonal fiber density.

Our current study investigated the brains of schizophrenia participants using the DBSI methodology. We hypothesized that due to the presumed psychopathology underlying SCZ, primary abnormalities found with this method would involve increased cellular diffusion fraction across multiple white matter tracts compared to healthy controls. In the study, we also included individuals with bipolar disorder, as an additional comparison group to establish specificity of findings. While diffusion imaging abnormalities have often been found in bipolar disorder, these are generally less extensive and severe than those seen in schizophrenia.^29^ We hypothesize that similarly, cellular diffusion fraction in BPD will be increased compared to the control group but substantially less than in SCZ participants.

## Methods

### Subjects

Imaging data were acquired from three participant groups, aged 18 to 30-year-old recruited through community advertisements and volunteer databases: 27 healthy young control (CON), 21 schizophrenia (SCZ), and 21 bipolar disorder (BPD). These participants were diagnosed on the basis of a trained research assistant who used the Structured Clinical Interview for DSM-IV Axis I Disorders (SCID-IV).^30^ CON subjects were required to have no lifetime history of psychotic or mood disorders. BPD participant patients were required to meet DSM-IV criteria for Bipolar I Disorder. To minimize clinical heterogeneity within the BPD group, only participants with a history of euphoric mania (versus mania characterized by primarily irritable mood) were included in the study. Written informed consent was obtained prior to participation, and all study protocols were approved by the Institutional Review Board at the Washington University School of Medicine in St. Louis, MO.

All participants were excluded if they: (a) met DSM-IV criteria for substance dependence or severe/moderate abuse during the prior 3 months; (b) had a clinically unstable or severe general medical disorder; or (c) had a history of head injury with documented neurological sequelae or loss of consciousness.

### Behavioral assessments

Recent symptoms (i.e., in the prior two weeks) were assessed using the Scale for the Assessment of Negative Symptoms (SANS), and the Scale for the Assessment of Positive Symptoms (SAPS).^31^ Chronic symptoms (i.e., prior year) were assessed using the Washington Early Recognition Center Affectivity and Psychosis (WERCAP) Screen, both affective (a-WERCAP) and psychosis (p-WERCAP) components.^32–34^

### Image acquisition

Structural T1w MRI images were acquired on a 3T Siemens Prisma with a 32 channel head coil using a 3D MPRAGE sequence^35^ (0.8 mm isotropic voxels, TR/TI = 2400/1000 ms, TE = 2.2 ms, flip angle = 8**°,** FOV = 256 ×240×166 mm, matrix size = 320 ×300, 208 sagittal slices, in-plane (iPAT) acceleration factor of 2). T2w volumes were also acquired at the same spatial resolution using the variable-flip-angle turbo-spin-echo 3D SPACE sequence^36^ (TR/TE=3200/564 ms; same FOV, matrix and in-plane acceleration). The diffusion MRI (dMRI) scans used the multi-band (MB) sequences from the Center for Magnetic Resonance Research, with 1.25 isotropic voxels, TR = 5000 ms, TE = 104 ms, 6/8 partial Fourier, and MB factor = 4. A full dMRI session included 6 runs (each approximately 8.5 min), representing 3 different gradient tables, with each table acquired once with anterior-to-posterior and posterior-to-anterior phase encoding polarities, respectively. Each gradient table includes approximately 90 diffusion weighting directions plus 6 b = 0 acquisitions interspersed throughout each run. Diffusion weighting consisted of 3 shells of b = 1000, 2000, and 3000 s/mm^2^ interspersed with an approximately equal number of acquisitions on each shell within each run.

### Image preprocessing

The diffusion data were preprocessed using the “DiffusionPreprocessing” stream of the HCPpipelines (v4.3.0),^37,38^ using the QuNex container (v0.91.11). This pipeline includes intensity normalization, susceptibility distortion correction (via FSL’s ‘topup’),^39^ and correction for eddy current distortions and motion via FSL’s ‘eddy’ tool.^40^ We used the advanced ‘eddy’ features of outlier replacement,^41^ slice-to-volume motion correction,^42^ and correction for susceptibility-by-movement interactions.^43^ The b-vectors were rotated to account for motion.^44^ Finally, the dMRI data was corrected for gradient nonlinearity distortion as part of resampling to the subject’s native T1w space from the HCP structural pipeline output (while maintaining the same 1.25 mm spatial resolution of the dMRI data). Following that preprocessing, two different diffusion models were implemented 1) diffusion tensor estimation using conventional FSL’s ‘DTI’ toolbox 2) modelled diffusion signal using DBSI and derived parameters such as cellular diffusion fraction, extra-axonal water, axonal fiber fraction, fiber axial, and fiber radial diffusivities. To perform voxel-wise analysis of all white matter tracts, Tract-based spatial statistics (TBSS) was used.^45^ DTI-derived FA images were calculated and projected onto the mean FA skeleton, which represents center of white matter tracts, and thresholded at 0.2. After pre-alignment and skeletonization stage, the resulting 4D data was used to perform voxel-wise statistics.

### DBSI

Diffusion data were modeled using the novel DBSI method. In DBSI, each of the potential pathological components, including inflammatory cell components, extra-cellular water/vasogenic edema, axonal injury/loss, and demyelination, is modelled within each voxel by a dedicated diffusion tensor. DBSI framework, as shown in Equation (1), considers diffusion-weighted MRI data as a linear combination of multiple anisotropic (crossing myelinated and unmyelinated axons of varied direction; the first term) and a spectrum of isotropic (cells, sub-cellular structure, and edematous water; the second term)^22^ diffusion tensors.

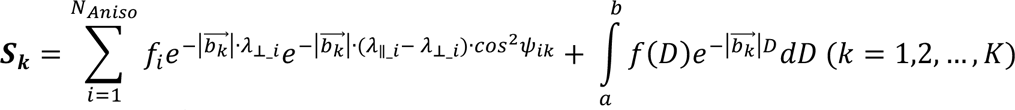

The quantities ***S***_***k***_ and 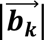 are the signal and b-value of the ***k^th^*** diffusion gradient, ***N***_***Aniso***_ is the number of anisotropic tensors (fiber tracts), ***Ψ***_***ik***_ is the angle between the ***k^th^***diffusion gradient and the principal direction of the ***i^th^*** anisotropic tensor, ***λ***_∥_***i***_ and ***λ***_⊥_***i***_ are the axial and radial diffusivities of the *i^th^* anisotropic tensor, ***f_i_*** is the signal intensity fraction for the ***i^th^***anisotropic tensor, and ***a*** and ***b*** are the low and high diffusivity limits for the isotropic diffusion spectrum ***f(D)***. In this study, the sum of ***f_i_*** is defined as the fiber fraction to reflect the total white matter axonal fiber density. The summation of the anisotropic diffusion component with radial diffusivity bigger than 0.1 (×10^−3^ mm^2^/s) is defined as the extra-axonal water fraction to reflect the extra-axonal water edema in WM. The summation of the isotropic diffusion components with apparent diffusion coefficient between 0 and 0.3 (×10^−3^ mm^2^/s) is defined as the cellular diffusion fraction to reflect the total cellularity in WM.

### Statistical analysis

Statistical analysis was performed using Kruskal-Wallis rank sum test for age and body mass index (BMI), and Pearson’s *χ*^2^ test for gender. These statistical analyses were performed with the software package R, Version 4.3.1 (https://www.r-project.org/).

To perform voxel-wise group comparisons, a permutation based nonparametric statistical test was performed using “randomise” − a part of FSL.^46^ Two sample non-parametric t-tests were performed between CON, BPD, and SCZ groups. The mean FA skeleton was used as mask and number of permutations were set to 5000. The significance threshold for group differences was set at *p* < 0.05. The statistical significance maps were corrected for multiple comparisons using threshold free cluster enhancement (TFCE) option in “randomise”. Similarly, group comparisons of DBSI metrics were performed.

For each DBSI measure, voxels with statistically significant difference (p < 0.05 after FWER correction) resulting from group wise comparisons 1) control vs schizophrenia, and 2) control vs bipolar disorder were added together, and binary mask was created. DBSI measures were summarized for each subject using corresponding mask. Finally, Pearson’s correlation was performed between DBSI measures (only with statistically significant group difference in randomized test) and clinical scores with the software package R. As age and gender were not statistically different across groups, we did not control for it in above analysis.

## Results

### Demographic and clinical profiles

**Table 1** shows demographic and clinical information across the participant groups. Mean age was similar across the three participant groups. The BPD group had substantially more females than males, while sex was more balanced in SCZ and CON.

**Table 1.** Baseline demographics and medication history across participant groups.

| Characteristic | CON, N = 27<sup>1</sup> | BP, N = 21<sup>1</sup> | SCZ, N = 21<sup>1</sup> | p-value<sup>2</sup> |
| --- | --- | --- | --- | --- |
| Age | 25.56 (3.3) | 27.00 (3.6) | 26.71 (4.1) | 0.3 |
| Gender |  |  |  | 0.076 |
| Female | 16 (59.3%) | 17 (81.0%) | 10 (47.6%) |  |
| Male | 11 (40.7%) | 4 (19.0%) | 11 (52.4%) |  |
| BMI | 26.47 (6.6) | 29.96 (7.4) | 29.69 (7.8) | 0.15 |
| BMI group |  |  |  |  |
| Underweight | 1 (3.7%) | 1 (4.8%) | 1 (4.8%) |  |
| Healthy | 12 (44.4%) | 4 (19.0%) | 5 (23.8%) |  |
| Overweight | 7 (25.9%) | 7 (33.3%) | 6 (28.6%) |  |
| Obese | 7 (25.9%) | 9 (42.9%) | 9 (42.9%) |  |
| Anti-psychotic medications |  |  |  |  |
| Typical neuroleptics | 0 | 0 | 3 (14.3%) |  |
| Atypical neuroleptics | 0 | 5 (23.8%) | 10 (47.6%) |  |
| Mood stabilizers |  |  |  |  |
| Lithium | 0 | 0 | 2 (9.5%) |  |
| Anti-convulsant | 0 | 6 (28.6%) | 2 (9.5%) |  |
| Both | 0 | 2 (9.5%) | 0 |  |
| Anti-depressants |  |  |  |  |
| SSRI/SNRI | 1 (3.7%) | 1 (4.8%) | 6 (28.6%) |  |
| Atypical | 0 | 2 (9.5%) | 2 (9.5%) |  |
| Both | 0 | 0 | 1 (4.8%) |  |
| Other/No medications |  |  |  |  |
| Other<sup>3</sup> | 0 | 3 (14.3%) | 3 (14.3%) |  |
| None<sup>4</sup> | 26 (96.3%) | 10 (47.6%) | 7 (33.3%) |  |

<sup>1</sup> Mean (SD); N (%)
<sup>2</sup> Kruskal-Wallis rank sum test; Pearson's Chi-squared test
<sup>3</sup> Participants with active benzodiazepines, anticholinergics, and beta-blockers.
<sup>4</sup> Participants with no active medication at all during the time of assessment.

### Voxel-wise analysis of DTI-derived fractional anisotropy, radial diffusivity, and axial diffusivity

**Table 2** shows white matter tracts where DBSI and DTI measures are significantly different in schizophrenia and bipolar disorder groups compared to control group.

**Table 2:** Summary of changes in DTI and DBSI measures across WM tracts and disease group. (Health Control=HC, Schizophrenia=SCZ, Bipolar disorder=BPD). ↑ denotes significant increase. ↓ denotes significant decrease and – denotes no statistically significant change. R denotes significant change in right hemisphere only, L denotes significant change in left hemisphere only, and arrow without R or L denotes change in both hemisphere.

| Diffusion parameter | DTI FA |  | DTI RD |  | DTI AD |  | Cellular diffusion fraction |  | Axonal fiber fraction |  | Extra-axonal water fraction |  | Fiber radial diffusivity |  | Fiber axial diffusivity |  |
| --- | --- | --- | --- | --- | --- | --- | --- | --- | --- | --- | --- | --- | --- | --- | --- | --- |
| Group comparison<br>WM tracts | HC vs. SCZ | HC vs. BPD | HC vs. SCZ | HC vs. BPD | HC vs. SCZ | HC vs. BPD | HC vs. SCZ | HC vs. BPD | HC vs. SCZ | HC vs. BPD | HC vs. SCZ | HC vs. BPD | HC vs. SCZ | HC vs. BPD | HC vs. SCZ | HC vs. BPD |
| Anterior thalamic radiation | ↓ | ↓ (L) | ↑ | -- | -- | -- | ↑ | -- | ↓ | -- | ↑ | ↑ | ↑ | ↑ | -- | -- |
| Cingulum (cingulate gyrus) | ↓ | ↓ | ↑ | -- | -- | -- | ↑ | -- | ↓ | -- | ↑ | ↑ | ↑ | ↑ | -- | -- |
| Cingulum (hippocampus) | ↓ | -- | ↑ | -- | -- | -- | ↑ | -- | ↓ | -- | ↑ | ↑ | ↑ | ↑ | -- | -- |
| Corticospinal tract | ↓ | -- | ↑ | -- | -- | -- | ↑ | -- | ↓ | -- | ↑ | ↑ | ↑ | ↑ | -- | -- |
| Forceps major | ↓ | -- | ↑ | -- | -- | -- | ↑ | -- | ↓ | -- | ↑ | ↑ | ↑ | ↑ | -- | -- |
| Forceps minor | ↓ | ↓ | -- | -- | -- | -- | ↑ | -- | ↓ | -- | ↑ | ↑ | ↑ | ↑ | -- | -- |
| Inferior fronto-occipital fasciculus | ↓ | ↓ (L) | ↑ | -- | -- | -- | ↑ | -- | ↓ | -- | ↑ | ↑ | ↑ | ↑ | -- | -- |
| Inferior longitudinal fasciculus | ↓ | -- | ↑ | -- | -- | -- | ↑ | -- | ↓ | -- | ↑ | ↑ | ↑ | ↑ (R) | -- | -- |
| Superior longitudinal fasciculus (temporal part) | ↓ | -- | ↑ | -- | -- | -- | ↑ | -- | ↓ | -- | ↑ | ↑ | ↑ | ↑ | -- | -- |
| Superior longitudinal fasciculus | ↓ | -- | ↑ | -- | -- | -- | ↑ | -- | ↓ | -- | ↑ | ↑ | ↑ | ↑ | -- | -- |
| Uncinate fasciculus | ↓ | ↓ (L) | -- | -- | -- | -- | ↑ | -- | ↓ | -- | ↑ | ↑ | ↑ | ↑ (L) | -- | -- |

As shown in **Figure 2A**, we observed low fractional anisotropy (FA) in SCZ compared to controls. The reduced FA was bilaterally distributed over anterior thalamic radiation, corticospinal tract, cingulum (both cingulate gyrus and hippocampus), forceps major, forceps minor, inferior fronto-occipital fasciculus, inferior longitudinal fasciculus, and superior longitudinal fasciculus. In contrast to findings in SCZ, the FA in BPD participants were minimal and were only observed unilaterally over anterior thalamic radiation, inferior fronto-occipital fasciculus, uncinate fasciculus, bilaterally in cingulum (cingulate gyrus) and forceps minor (**Figure 2B**).

**Figure 2:**
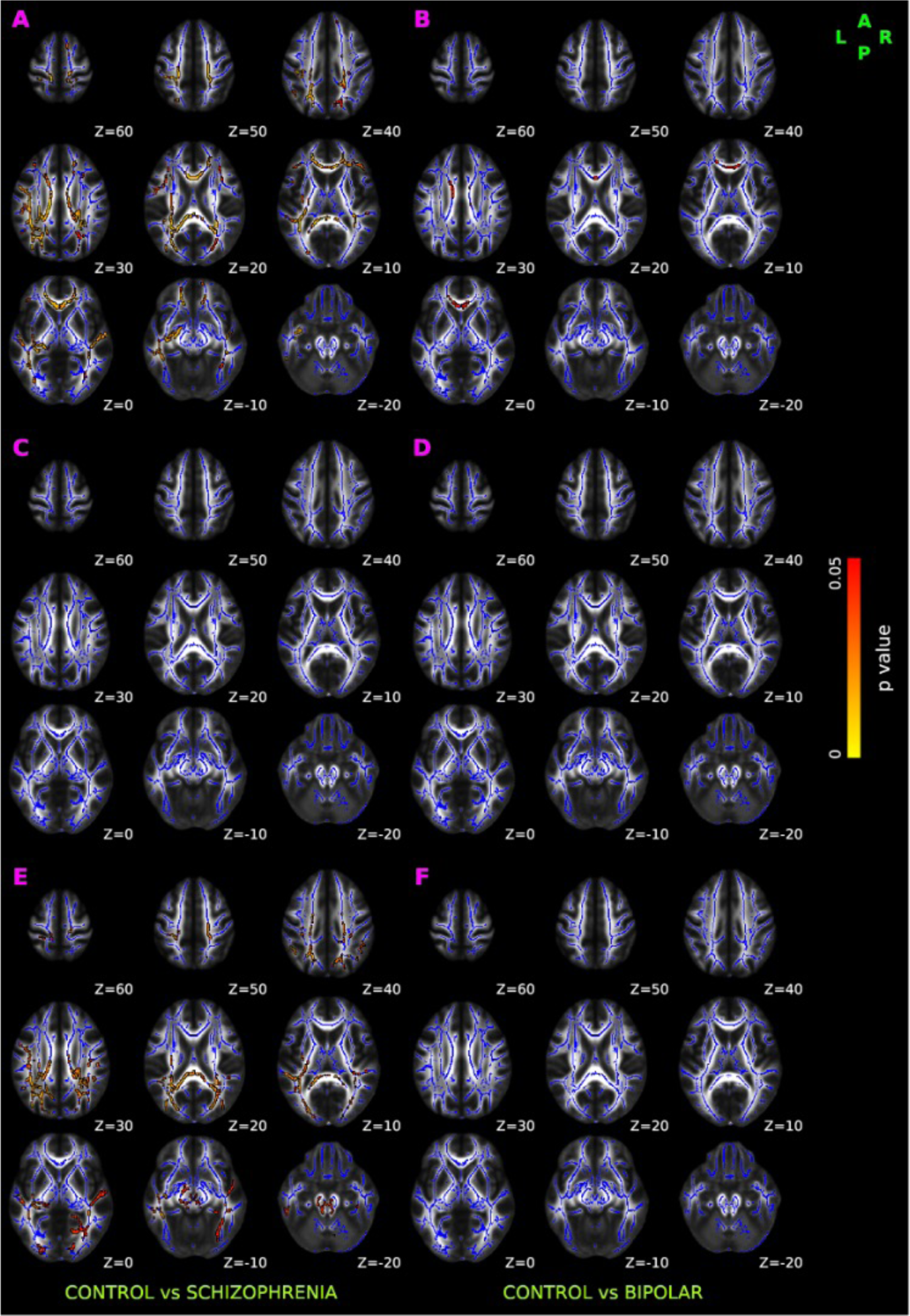
TBSS results of DTI-derived fractional anisotropy (FA), axial diffusivity (AD), and radial diffusivity (RD). Red-yellow overlay represents statistically significant difference in FA (Panel A and Panel B), AD (Panel C &D), and RD (Panel E & F) in schizophrenia and bipolar disorder group relative to normal control group (p < 0.05, FWER correction by TFCE). To aid visualization, regions showing changes in DTI metric are dilated using 3×3×1 kernel in FSL. Results are shown overlaid on the MNI 152-T1 1mm template and the mean FA skeleton (Blue).

No significant axial diffusivity (AD) abnormalities were seen in either SCZ (**Figure 2C**) or BPD (**Figure 2D**) participants. SCZ subjects however had significantly higher radial diffusivity (RD) than controls in similar regions where decreased FA was observed, as shown in **Figure 2E**. No significant abnormalities in RD were observed in BPD (**Figure 2F**).

### Voxel-wise analysis of cellular diffusion fraction

Relative to controls, SCZ had significantly increased cellular diffusion fraction — a putative marker of white matter inflammation (**Figure 3**A), bilaterally spread over anterior thalamic radiation, corticospinal tract, cingulum bundle, forceps major, forceps minor, inferior fronto-occipital fasciculus, uncinate fasciculus, and superior longitudinal fasciculus.

**Figure 3.**
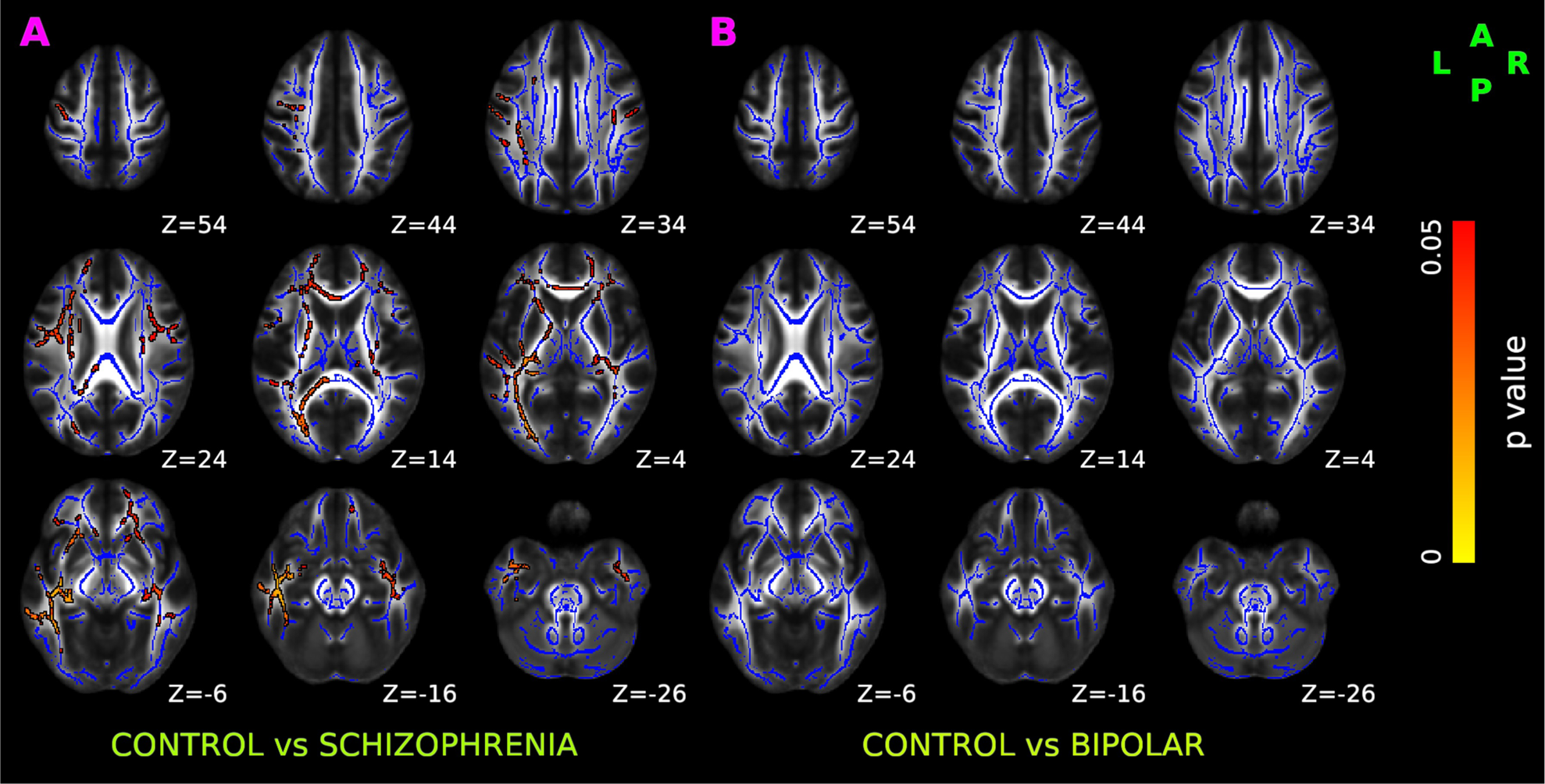
TBSS results of cellular diffusion fraction in healthy control, schizophrenia, and bipolar disorder participants. Red-yellow overlay represents regions with statistically significant changes in cellularity ratio in schizophrenia and bipolar disorder patients (Panel A & B respectively) compared to control subjects (p < 0.05, FWER correction by TFCE). To aid visualization, regions showing increased cellularity ratio are dilated using 3×3×1 kernel in FSL. Results are shown overlaid on the MNI 152-T1 1mm template and the mean FA skeleton (Blue).

No significant alterations in cellular diffusion fraction were observed in BPD after FWER correction (**Figure 3**B).

### Voxel-wise analysis of axonal fiber fraction

Compared to controls, SCZ had statistically decreased fiber fraction (**Figure 4**A) bilaterally in anterior thalamic radiation, corticospinal tract, cingulum bundle, forceps major, forceps minor, inferior fronto-occipital fasciculus, uncinate fasciculus, and temporal region of superior longitudinal fasciculus.

**Figure 4.**
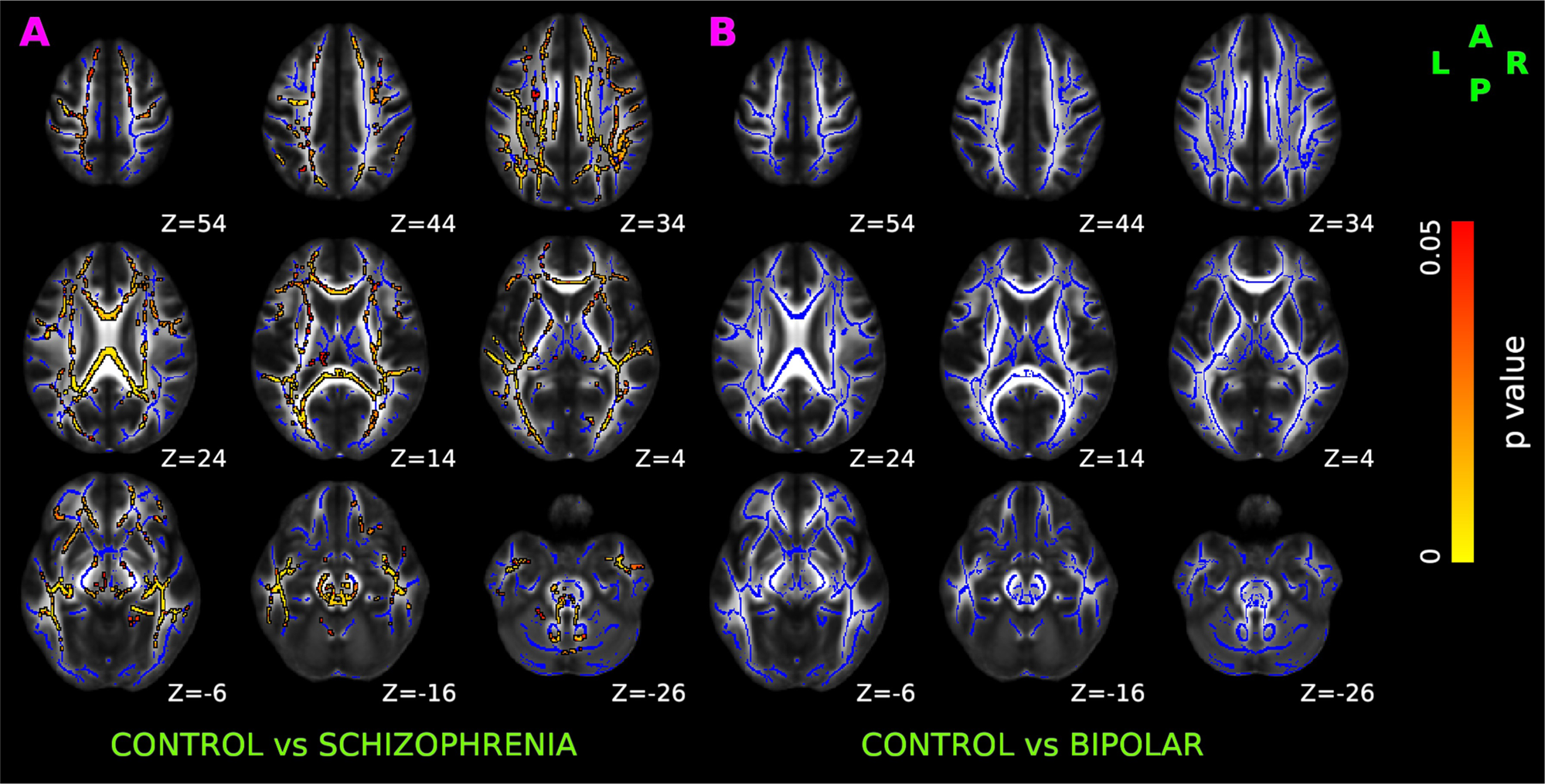
TBSS results of axonal fiber fraction in healthy control, schizophrenia, and bipolar disorder participants. Red-yellow overlay represents statistically significant changes in neuronal density (fiber ratio) in schizophrenia and bipolar disorder subjects (Panel A & B) relative to controls (p < 0.05, FWER correction by TFCE). To aid visualization, regions showing decreased fiber1 ratio are dilated using 3×3×1 kernel in FSL. Results are shown overlaid on the MNI 152-T1 1mm template and the mean FA skeleton (Blue).

Compared to controls, changes in the axonal fiber fraction were not statistically significant in BPD (**Figure 4**B).

### Voxel-wise analysis of extra-axonal water fraction

As shown in **Figure 5**, compared to controls, SCZ and BPD had statistically increased extra-axonal water fraction (Panel A and Panel B respectively). In both SCZ and BPD, increased extra-axonal water fraction was present bilaterally over anterior thalamic radiation, corticospinal tract, cingulum bundle, forceps major, forceps minor, inferior fronto-occipital fasciculus, inferior longitudinal fasciculus, uncinate fasciculus, and superior longitudinal fasciculus.

**Figure 5.**
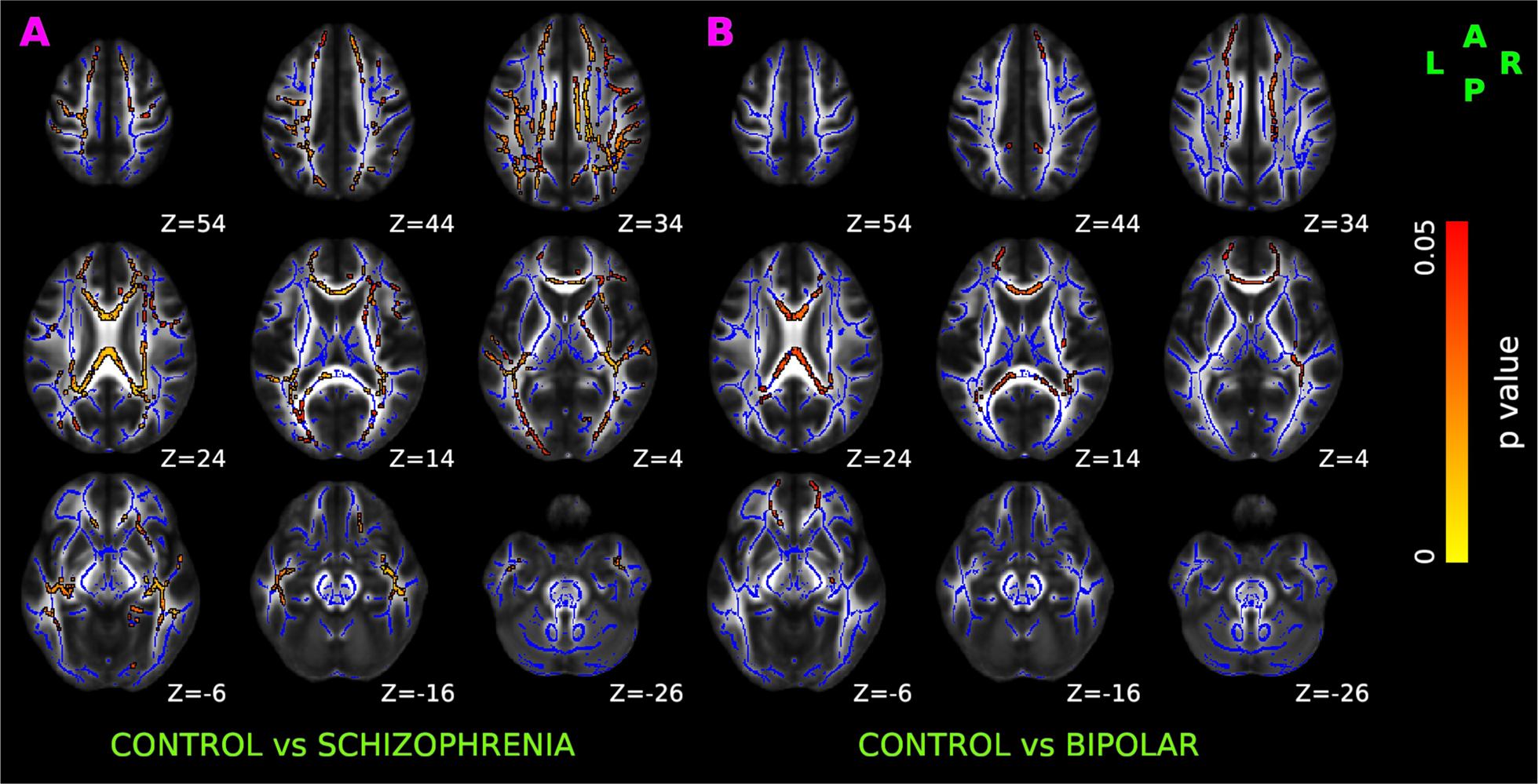
TBSS results of DBSI-derived extra-axonal water in healthy control, schizophrenia, and bipolar disorder participants. Red-yellow overlay represents a statistically significantly increased extra-axonal water in schizophrenia and bipolar disorder patients (Panel A & B respectively) relative to controls (p < 0.05, FWER correction by TFCE). To aid visualization, regions showing increased fiber23 ratio are dilated using 3×3×1 kernel in FSL. Results are shown overlaid on the MNI 152-T1 1mm template and the mean FA skeleton (Blue).

Group comparison between SCZ and BPD did not reveal significant differences in the extra-axonal water fraction.

### Voxel-wise analysis of DBSI-derived fiber axial and fiber radial diffusivity

Compared to controls, fiber axial diffusivity changes in SCZ (**Figure 6**A), and BPD (**Figure 6**B) did not survive the significance threshold after FWER correction.

**Figure 6.**
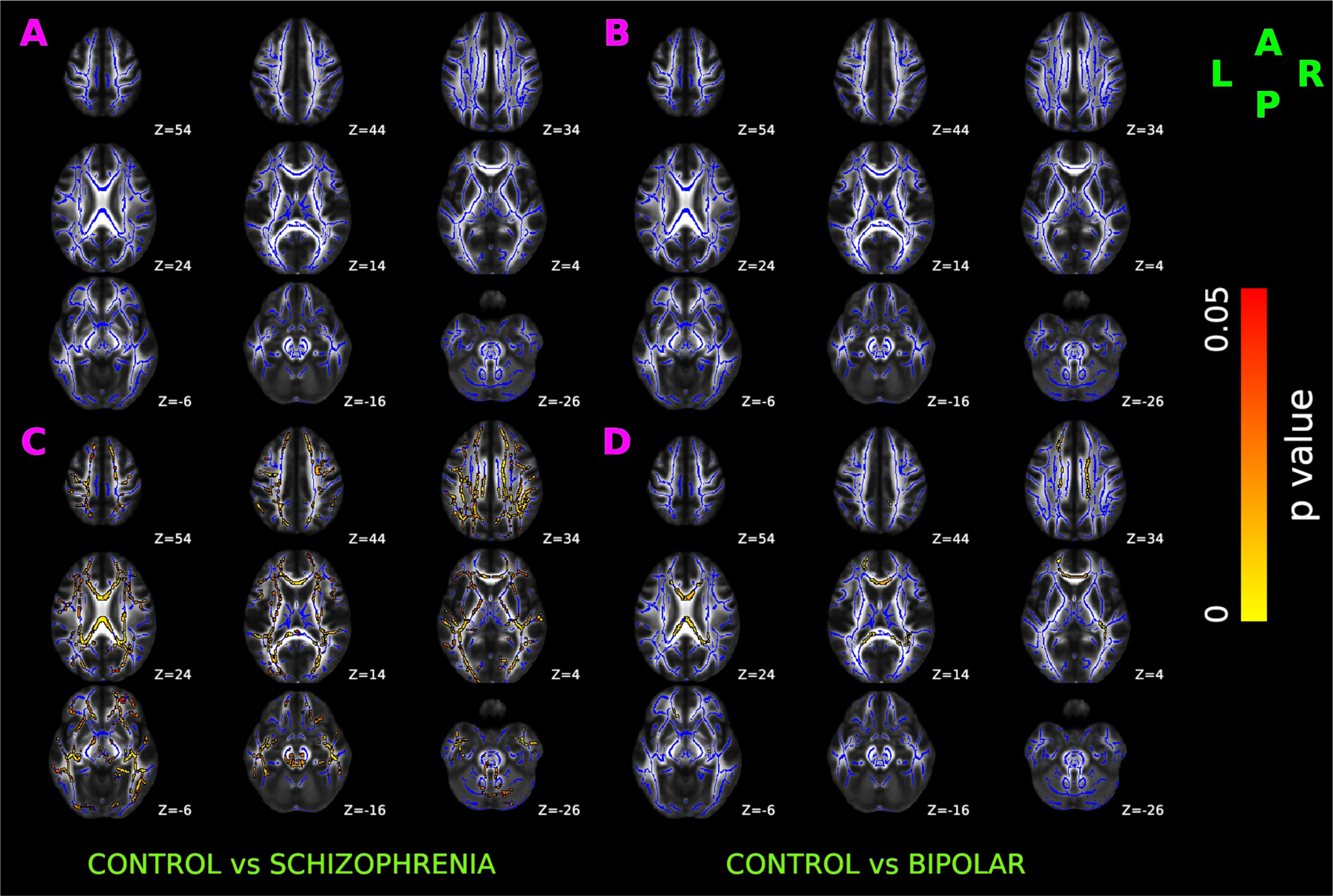
TBSS results of DBSI-derived axial and radial diffusivity in healthy control, schizophrenia, and bipolar disorder participants. Red-yellow overlay represents statistically significant difference in fiber axial diffusivity (Panel A & B) and fiber radial diffusivity (Panel C & D) in schizophrenia and bipolar disorder group (respectively) relative to controls (p < 0.05, FWER correction by TFCE). To aid visualization, regions showing increased radial diffusivity are dilated using 3×3×1 kernel in FSL. Results are shown overlaid on the MNI 152-T1 1mm template and the mean FA skeleton (Blue).

Relative to controls, SCZ (**Figure 6**C) had significantly increased fiber radial diffusivity, spread bilaterally over anterior thalamic radiation, corticospinal tract, cingulum bundle, forceps major, forceps minor, inferior fronto-occipital fasciculus, inferior longitudinal fasciculus, uncinate fasciculus, and superior longitudinal fasciculus. In BPD (**Figure 6**D), increased fiber radial diffusivity was observed bilaterally over anterior thalamic radiation, corticospinal tract, cingulum bundle, forceps major, forceps minor, inferior fronto-occipital fasciculus, superior longitudinal fasciculus, unilaterally in right inferior longitudinal fasciculus and left uncinate fasciculus.

### Correlation analysis between WERCAP symptom score and DBSI measures

Across healthy control and SCZ, a correlation of psychotic symptom scores against DBSI measures partialled for diagnosis showed significant effects (r>0.55; p<0.0001) for cellular diffusion fraction (**Figure 7A**), fiber radial diffusivity (**Figure 7B**), fiber fraction (**Figure 7C**) and extra-axonal water fraction (**Figure 7D**). The relationship between psychotic symptoms and DBSI metrics separately in SCZ and control participants did not meet statistical significance.

**Figure 7.**
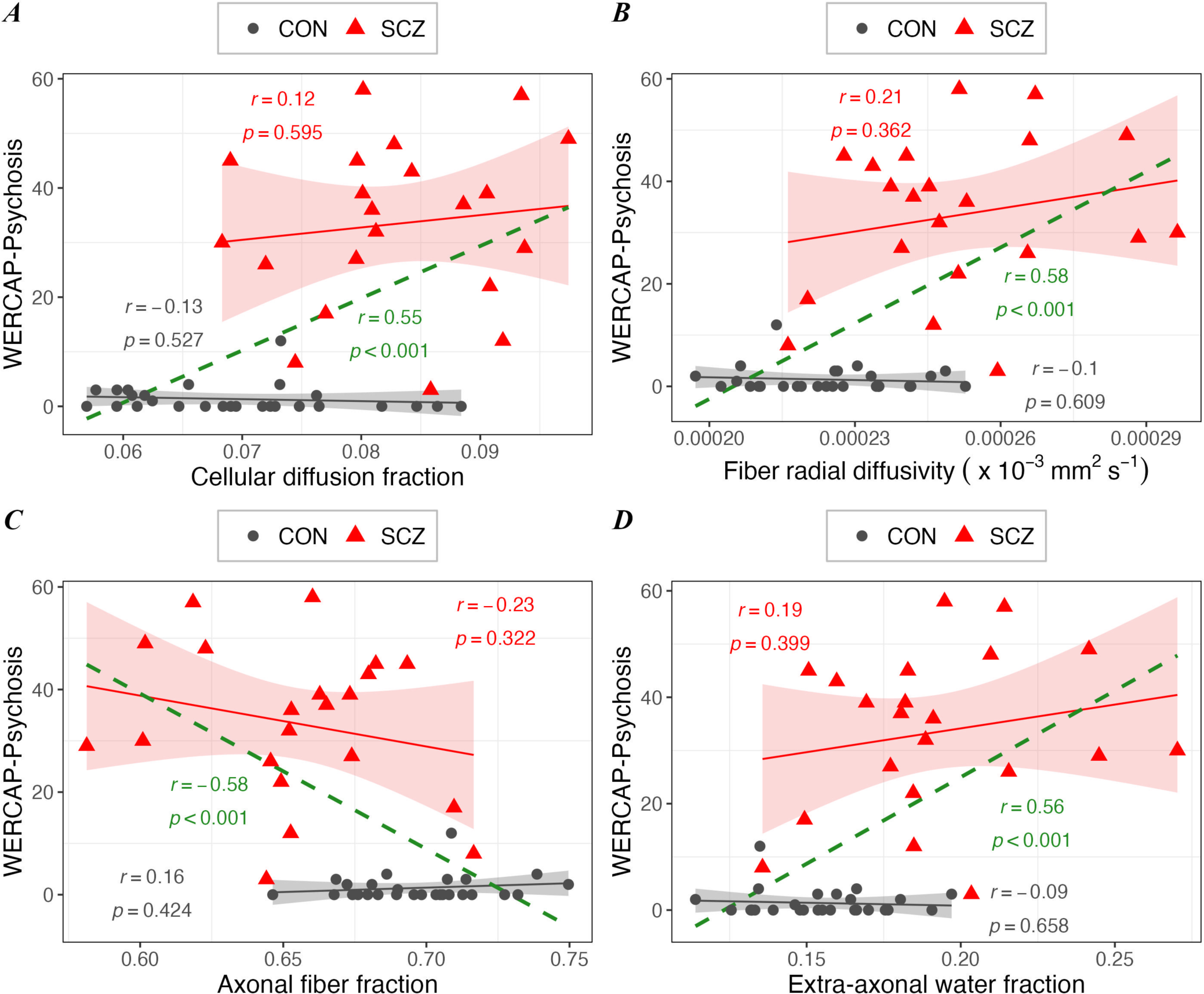
Relationship between WERCAP-Psychosis symptom score and DBSI measures in schizophrenia (in red), in control (in black), and in combined group (in green). Scatterplots showing significant correlation between WERCAP-Psychosis score and DBSI measures – (A) Cellular diffusion fraction, (B) DBSI-derived radial diffusivity, (C) Fiber density, and (D) extra-axonal water. Shaded region surrounding regression lines shows 95% confidence interval.

Similar findings were observed for the relationship between mood symptom scores and DBSI measures (**Figures 8A-D**), though the relationship was weaker (r>0.28; p<0.05).

**Figure 8.**
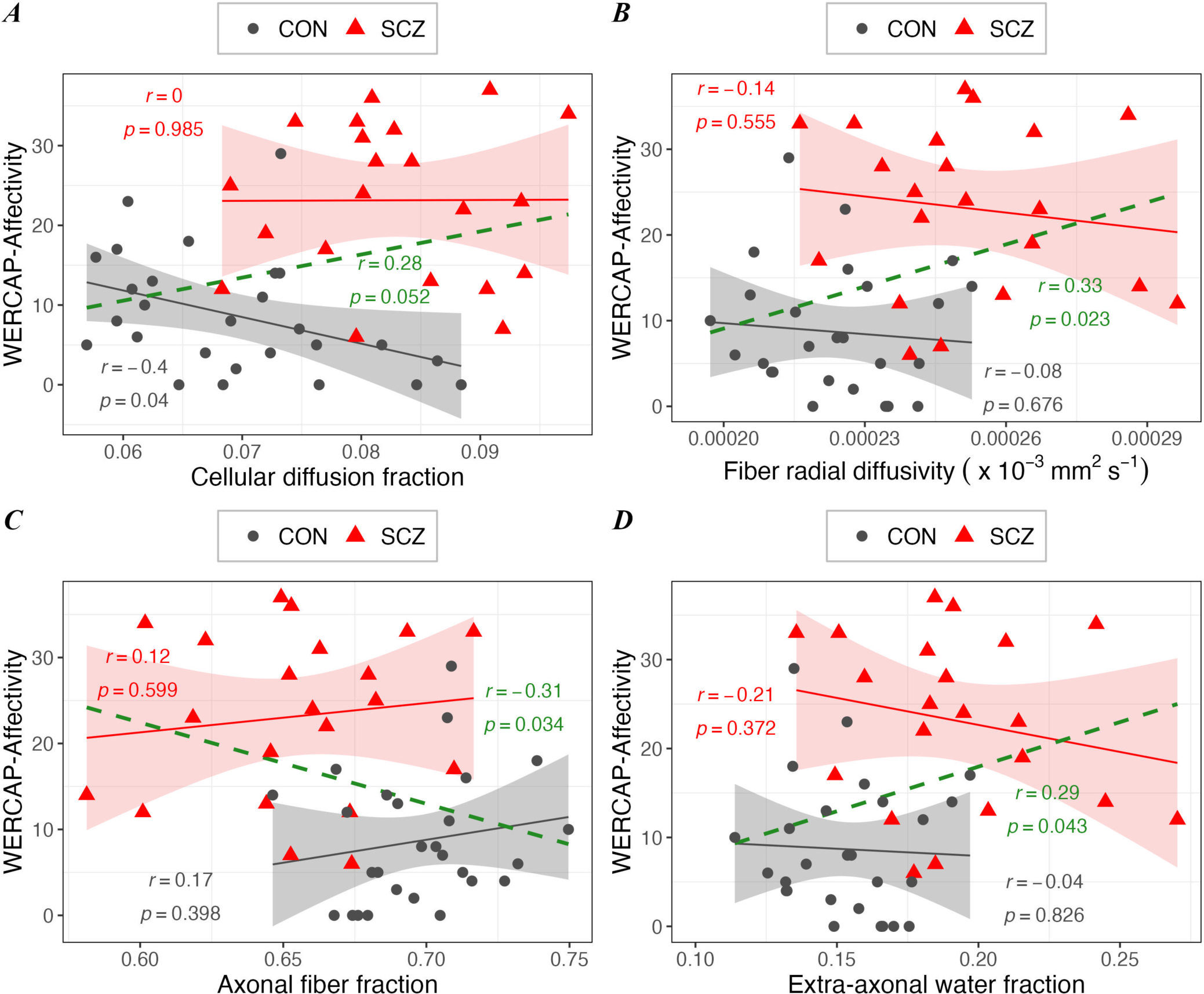
The associations between WERCAP-Affectivity symptom score and DBSI measures in schizophrenia (in red), in control (in black), and in combined group (in green). Scatterplots showing significant correlation between WERCAP-Affectivity score and DBSI measures – (A) Cellular diffusion fraction, (B) DBSI-derived radial diffusivity, (C) Fiber density, and (D) extra-axonal water. Shaded region surrounding regression lines shows 95% confidence interval.

## Discussion

Our study expands the novel neuroimaging, diffusion basis spectrum imaging (DBSI), for the first time to the study of individuals with schizophrenia and bipolar disorder. DBSI is particularly promising in the study of those with psychiatric disorders, which generally do not have validated biomarkers with clinical utility. This method has been shown to distinguish various pathophysiologic changes of brain white matter in vivo in postmortem studies of central nervous system disorders.^47^

The commonly used diffusion imaging metrics DTI-derived FA, RD and AD have been valuable in understanding brain microstructural abnormality in disease. However, establishing underlying when these metrics are abnormal is complicated by the fact that different pathologies can contribute to it. For example, while high RD is suggested to be largely related to axonal demyelination, other factors like edema and cellularity can confound symptoms. By delineating individual white matter components, DBSI-derived RD and AD are posited to minimize confounding effects, compared to DTI-derived metrics. Our study found a generalized bilateral increase in white matter RD in schizophrenia, without associated abnormality in AD. This suggests that demyelination and not axonal degeneration is likely the underlying microstructural pathology.^48^ A minimally elevated RD was also seen in BPD participants but only with DBSI, not with conventional DTI. Demyelination of white matter tracts is consistent with previous studies of schizophrenia. Postmortem samples have shown oligodendrocytes have been found to show consistent signs of dystrophia, apoptosis and/or necrosis, and their numerical density have been significantly reduced in schizophrenia.^49^ Schizophrenia brain samples have also been characterized by downregulation of myelin-specific structural proteins and a reduction in mRNA levels of oligodendroglial lineage transcription factors, such as OLIG1, OLIG2 and SOX10.^50^ Myelin-related abnormalities in schizophrenia would lead to disconnection of neural networks by impairing the saltatory conduction and information conduction between neurons.^51^ Findings of high RD in schizophrenia appear to be disorder-specific, as similar changes were seen to a milder degree, consistent with fewer white matter abnormalities found in other studies.^52^

A notable finding in our study was an elevated cellular diffusion fraction across multiple white matter regions exclusively in schizophrenia subjects. The DBSI metric generally correlates with elevated inflammatory cells and is seen in brain conditions with inflammatory underpinnings, such as multiple sclerosis,^22^ HIV infection,^28^ and autoimmune encephalitis in mice.^53^ There has been increasing evidence for an inflammatory signature in the brain in schizophrenia, including overexpression of pro-inflammatory cytokines,^54^ changes to the blood-brain barrier with macrophage infiltration^55^ and possible microglia activation.^56^ Schizophrenia has also been associated with elevated peripheral inflammatory markers,^14,15^ binding of pro-inflammatory PET tracers,^12^ and genetic risk factors of immune dysregulation.^18^ Inflammatory markers have been found to precede the onset of illness^13,16^ suggesting a progressive pathological process leading to an onset of schizophrenia. The absence of cellular diffusion fraction abnormalities in our bipolar disorder population suggests the specificity of this DBSI marker to schizophrenia. Although not clear from our results, it is worth speculating whether increased cellular diffusion fraction is confounded by increased density of interstitial white matter neurons in schizophrenia,^57^ reflecting a deficiency in interneuron migration from white matter to cortex during development. Migration of these subplate neurons has been associated with environmental insults including maternal infection in early life and cannabis use during childhood and adolescence.^58^

Another major finding in our study involved a decreased fiber fraction in schizophrenia participants, a marker of apparent axonal fiber density. Here again, findings were bilateral and generalized to multiple white matter tracts throughout the brain. There have been few studies investigating white matter fiber density in schizophrenia, however, results have been somewhat consistent. A ‘fixel-based’ analysis of diffusion imaging data indicated lower primary fiber density in the postcentral and posterior corpus callosum in schizophrenia patients,^59^ although with higher associated fiber density in secondary and tertiary fibers. A diffusion imaging study of first-episode psychosis using NODDI analysis found decreased neurite density in multiple white matter tracts bilaterally or on the left^60^ suggesting these changes occur early in the course of illness. Aberrant white matter neurodevelopment is likely to precede the onset of schizophrenia, suggesting structural vulnerability to the disorder.^61^ Reduced fiber density might arise from aberrant neuronal migration, axon guidance, or synaptic pruning,^62^ and underlie much of the dysconnectivity characteristic of the disorder. Future longitudinal studies of fiber density in those at clinical high risk for psychosis (CHR-P) using DBSI would provide further insights into changes in white matter structure with psychosis progression. In contrast to fiber density, it is notable that postmortem studies of neuronal density in schizophrenia have been mixed, with most postmortem studies reporting both increased^63^ and decreased^59^ neuronal density in schizophrenia, along with decreased dendritic spine density^64^ believed to underlie altered synaptic connectivity and clinical presentation of schizophrenia patients.

Along with increased cellular diffusion fraction and decreased fiber fraction, we also found elevated extra-axonal water fraction, a marker of extra-axonal water edema, across multiple white matter tracts in schizophrenia participants. Taken together, these findings suggest that in our schizophrenia patients, there exists a pattern of inflammatory demyelination of low-density fiber tracts, with associated extra-axonal water edema. In bipolar disorder, microstructural changes suggested by DBSI results were milder degrees of demyelination and extra-axonal water edema only. Our findings also suggest that increased cellularity and decreased fiber density were specific to schizophrenia participants and may thus represent a disorder-specific biomarker representing a more extensive dysconnectivity syndrome. More importantly, it suggests that schizophrenia involves aberrant neurodevelopment of white matter tracts.

There are some limitations to our study. Due to the relatively small sample size, we did not control for potential confounders such as medications and substance use which are more common in those with severe psychiatric illnesses than in healthy controls. Antipsychotic medications for example have been associated with reductions in white matter FA^65^ and increases in white matter volume.^66^ Similarly, earlier age of first cannabis use has been linked to decreased FA and increased RD in long-range tracts.^67^ Thus, some of the white matter abnormalities observed in this study may be related to above factors, requiring larger studies to investigate. However, the absence of substantial findings in those with bipolar disorder, which has among the highest rates of substance use may partly argue against a major role in our findings,^68^ patterns of substance use tend to differ between disorders, with schizophrenia subjects more likely using non-alcoholic drugs.^68^ Secondly, while DBSI has been validated in preclinical models of white matter injury^69^ and in autopsied multiple sclerosis brain specimens,^70^ these results may not be completely valid in schizophrenia patients. A validation study involving postmortem schizophrenia samples is currently ongoing by our group. Thirdly, while the characteristic DBSI findings reported were observed in the majority of schizophrenia participants, they were not present in several participants, consistent with the heterogeneous brain profiles of the disorder.^7^ Larger sample sizes and longitudinal investigations would provide deeper insight into distinct schizophrenia biotypes, and brain changes at varying time points across the life span.

Our study shows that a novel diffusion imaging method, DBSI, could be used to safely identify unique microstructural changes in the brain, and has potential for clinical application in the future, including personalizing therapies. Specific white matter structural abnormalities, such as decreased neurite density, likely relate to poorer long-term prognosis and response to specific medications,^60^ and can therefore aid in the treatment selection. DBSI metrics could be used to monitor treatment effects, and may also identify individuals most at risk for developing psychosis who would benefit from preventative interventions before the onset of psychosis.

## Acknowledgments and Disclosures

This work was supported by the National Institute of Mental Health (Grant No. R01MH104414 [to DM]; and Grant No. R21MH131962 [to DM and YW];). All authors report no biomedical financial interests or potential conflicts of interest.

